# Modelling the effects of cerebral microthrombi on tissue oxygenation and cell death

**DOI:** 10.1101/2021.01.16.426717

**Authors:** Yidan Xue, Wahbi K. El-Bouri, Tamás I. Józsa, Stephen J. Payne

## Abstract

Thrombectomy, the mechanical removal of a clot, is the most common way to treat ischaemic stroke with large vessel occlusions. However, perfusion cannot always be restored after such an intervention. It has been hypothesised that the absence of reperfusion is due to the clot fragments that block the downstream vessels. In this paper, we present a new way of quantifying the effects of cerebral microthrombi on oxygen transport to tissue in terms of hypoxia and ischaemia. The oxygen transport was simulated with the Green’s function method on physiologically accurate microvascular cubes, which was found independent of both microvascular geometry and length scale. The microthrombi occlusions were then simulated in the microvasculature, which were extravasated over time with a new vessel extravasation model. The tissue hypoxic fraction was fitted as a sigmoidal function of vessel blockage fraction, which was then taken to be a function of time after the formation of microthrombi occlusions. A novel hypoxia-based 3-state cell death model was finally proposed to simulate the hypoxic tissue damage over time. Using the cell death model, the impact of a certain degree of microthrombi occlusions on tissue viability and microinfarct volume can be predicted over time. Quantifying the impact of microthrombi on oxygen transport and tissue death will play an important role in full brain models of ischaemic stroke and thrombectomy.

## 1 Introduction

Stroke is one of the leading causes of death and disability in the world, while ischaemic stroke accounts for about 85% of cases^1^. During ischaemic stroke, large vessel occlusions lead to a significant reduction in cerebral blood flow (CBF) to regions of the brain and hence to brain tissue death^2^. Thrombectomy, the mechanical removal of a clot, is the most common surgical treatment to recanalize the vessel^3^. It remains unclear why complete reperfusion cannot always be achieved after mechanical recanalization. It has been hypothesised that clot fragments are formed during the intervention and that the resulting microthrombi block the downstream microvasculature^4–6^.

Due to the limited resolution of current imaging techniques, it may not be possible to monitor microthrombi occlusions inside the human cerebral microvasculature, and hence to study the related clinical outcomes^5,7,8^. Rodent models have thus been widely used to investigate cerebral micro-embolisms and micro-infarcts. However these studies depend significantly on the assumed similarity between human and rodent cerebral vascular geometry^9–12^. *In silico* modelling can thus be used as an alternative to study the effects of microthrombi on human brain tissues. Previous *in silico* models of the microvasculature include models of the capillary beds^13–15^ and penetrating vessels^14,16^ generated from morphological data of the human cortex^17–19^. In one recent study, the effects of a penetrating vessel occlusion were simulated for the first time at a length scale comparable to that of MRI voxels, which can be directly validated against clinical images^20^.

The INSIST (IN-Silico trials for treatment of acute Ischemic STroke, www.insist-h2020.eu)^21^ is a multi-disciplinary project, which aims to build an *in silico* platform with virtual patients and computational models. The platform can be used to evaluate medical interventions and devices for the treatment of ischaemic stroke. As part of the project, we are developing organ-scale brain models to simulate the blood flow, oxygen transport and infarct progression during an ischaemic stroke^22,23^. A model that simulates clot fragmentation and the resulting effects on perfusion and oxygen transport can be directly coupled with the current whole brain model to investigate the reasons for reperfusion failure after thrombectomy.

There have been some *in silico* models proposed to determine the impact of microthrombi occlusions on perfusion in the microvasculature^24–26^. In our previous study^26^, we studied the effects of clot fragmentation on perfusion after thrombectomy. The clot fragmentation and micro-emboli shower simulations were based on *in vitro* experimental data^6^. Blood flow was modelled inside our statistically accurate microvasculature models, including penetrating arterioles and capillaries^15,16,20^. The perfusion drops in microvascular voxels and cortical columns and their relationships with different blockage percentage were investigated. However, none of the aforementioned models simulate the impact of microthrombi on tissue oxygenation and cell death. This will be a necessary step in order to validate our *in silico* whole brain model against clinical imaging data from post-thrombectomy patients.

In this paper, we thus present the effects of micro-occlusions on the oxygen transport and tissue oxygenation to provide a direct link between these occlusions and the response of the tissue. We do this first using existing Green’s function methods^27^. The microvascular recovery^12,28,29^ is then simulated by a new extravasation model. Additionally, a novel and validated hypoxia-based cell death model is proposed to predict the hypoxic tissue damage over time. These new models will play a key role in our organ scale *in silico* brain modelling and future validation against clinical data.

## 2 Methods

### 2.1 Capillary networks

The periodic, statistically accurate human capillary networks^15^ used here were generated in 3D using a statistical algorithm^13^ (Fig. 1a). The artificial networks match the morphometric information available on human cerebral capillary networks, which includes connectivity, vessel density, and the histograms of vessel length and diameter^17^. Given both flow and spatial periodicity, these network cubes can easily be connected to represent a larger domain of arbitrary size human capillary networks. It should be noted that the capillary networks can also be homogenised^30^. Hence the capillary bed can be treated as a porous medium with a permeability tensor representing the ease through which blood flows through the network.

**Figure 1.**
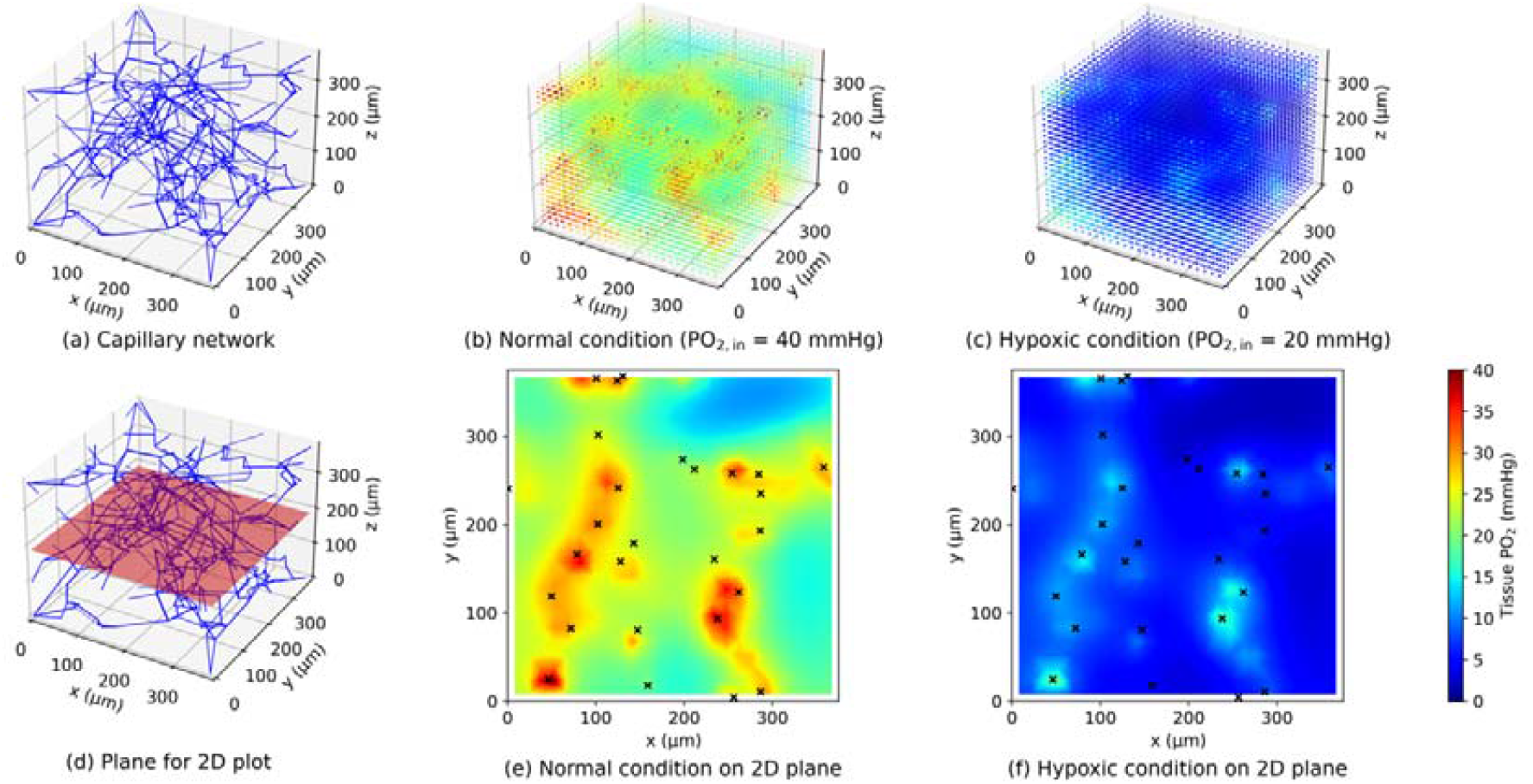
3D visualisation of an example capillary network (a) and the tissue oxygenation inside it solved by the Green’s function method under normal condition (b) and hypoxic condition (c). A horizontal plane (d) is placed in the centre of the capillary network and the tissue oxygenation on the plane is plotted under both normal (e) and hypoxic (f) conditions. The black crosses indicate the intersecting points between the centrelines of blood vessels and the plane.

In order to demonstrate the independence of oxygen transport results on length scales, two sizes of capillary cubes were selected: 375 μm and 625 μm. These two sizes are both within the convergence range of the permeability tensor, which indicates their similar geometrical characteristics in the perfusion calculation^15^. At each size, 10 cubes were used to investigate the dependence of oxygen transport on the capillary network geometry: this is required as the cubes match human data on a statistical basis, hence multiple cubes must be considered to obtain statistically accurate converged results. One example 375-micron cube is shown in Figure 1a.

### 2.2 Blood flow simulation

The blood flow calculations were conducted in MATLAB R2019b (MathWorks, USA). The same method was used here as in previous studies^15^ to calculate the blood flow in each vessel. Poiseuille flow and periodic boundary conditions were again assumed. The baseline value of perfusion in each cube was scaled to 55 ml/100g/min under normal conditions, by adjusting the pressure difference, where perfusion was calculated as the ratio of the inlet volumetric blood flow rate and the cube volume divided by tissue density (taken here to be 1.05 g/cm^3^).^31,32^ The key equations and assumptions of the blood flow simulation are summarised in Appendix A.

### 2.3 Oxygen transport simulation

The oxygen delivery from microvasculature to tissue was simulated using the well-established Green’s function method^27,33^. We used the publicly available implementation, written in C++^1^. The method has recently been applied to oxygen delivery^34,35^, and we therefore use this implementation. The cerebral metabolic rate of oxygen (CMRO_2_) was assumed to follow a Michaelis-Menten relationship dependent upon partial pressure of oxygen (PO_2_). The relationship between the blood oxygen saturation and the blood PO_2_ was assumed to be described by a Hill equation. The governing equations of these models are summarised in Appendix B. All of the model parameters remained the same as previously used^27^, except for the maximum CMRO_2_, which was set as 6.72×10-^4^ cm^3^/(cm^3^ · s) based on human data^36^ and the haematocrit was assumed to be a constant value of 0.45 which has been used previously to simulate the blood flow^15^.

The inlet PO_2_ (PO_2, in_) was then varied from 13 mmHg to 50 mmHg in steps of 1 mmHg in each capillary cube, in order to explore a wide range of values. After running the simulation, PO_2_ and oxygen consumption at each tissue point were analysed to obtain the average tissue PO_2_, hypoxic fraction (taken here to be the fraction of tissue at a value of PO_2_ below 10 mmHg), CMRO_2_ and oxygen extraction fraction (OEF), which is the fraction of blood oxygen taken by the tissue. The averaged values of all tissue points were used to calculate the CMRO2 and OEF inside the capillary cubes. The pre-processing and post-processing of the results were performed using custom Python scripts. An example capillary cube and its Green’s function solutions under both normal and hypoxic conditions are shown in Fig. 1.

### 2.4 Simulation of micro-thrombi inside the microvasculature

To mimic the presence of micro-thrombi inside the microvasculature, random occlusions were introduced into a 375-micron capillary network by using Python’s ‘random’ function to select the target vessel. A specified number of vessels was occluded to give a blockage fraction ranging from 1% to 20% in steps of 1% (rounded to the closest integer). At each blockage fraction, 5 different simulations were carried out with PO_2, in_ set to a value of 40 mmHg.

It should be noted that in order to maintain the periodicity of the network, the vessels connected to the boundary nodes were excluded from occlusions. In addition, any occlusion simulation that divided the network into multiple isolated parts was removed from consideration in order to keep the connectivity, so that the conductance matrix (Eq. A2) remained invertible and the flow could be solved with the existing algorithm^15^. For the same purpose, the maximum blockage fraction was set as 20%. After randomly blocking the vessels, the same flow and Green’s function simulation and data post-processing were performed as described earlier.

### 2.5 Vessel extravasation simulation

After the initial blockage, the blockage fraction was assumed to decay exponentially with time due to thrombus extravasation^28,29^. The blockage fraction thus can be written as

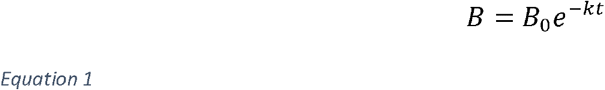

where *B*_0_ is the initial blockage fraction and *k* is the time constant. A recent extravasation study in a rat model^12^ was used for fitting the parameter *k* with the curve fitting function (non-linear least square optimization) built in SciPy^2^, an open-sourced Python library, as shown in Figure 2. The value of *k* in Eq.1 was found to be 2.03 × 10^−6^ in the unit of 1/day, when 4 data points, at 0, 1, 7, 28 days, were used for fitting. It should be noted that the extravasation model here is fitted with animal data (a rat model) due to the lack of any available human data. We thus assume the extravasation processes in rat brain tissue and human brain tissue to be very similar^29^, although it would be straightforward to adapt this model for human brain tissue should additional data become available.

**Figure 2.**
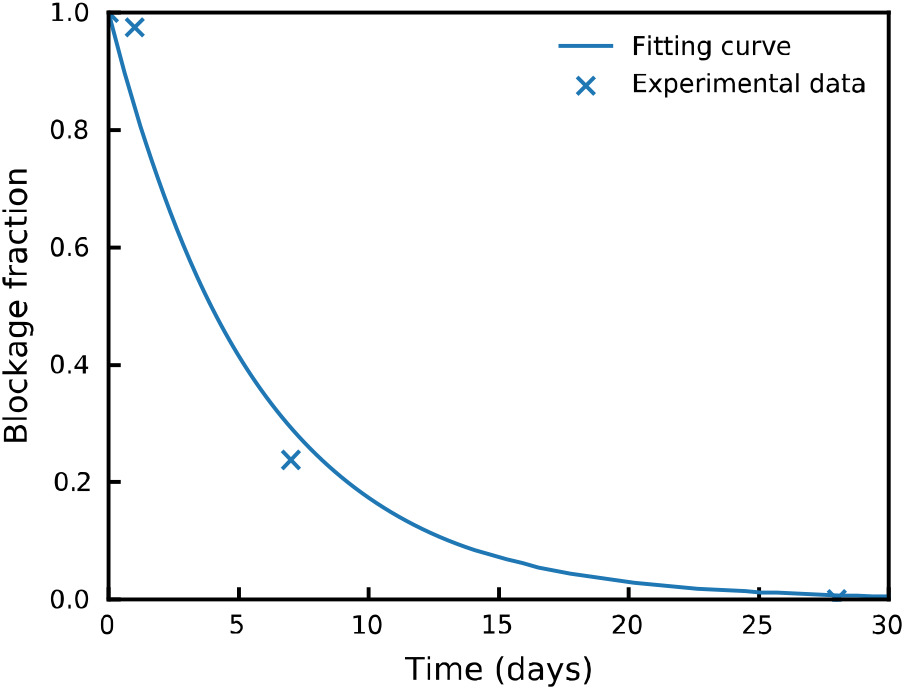
The extravasation of occluded vessels over time. The experimental data^12^ are taken from a rat model.

### 2.6 Three-state cell death model

A new hypoxia-based 3-state cell death model is proposed here based on the concept from previous work studying hyperthermic cell death^37^. Although this is a significant simplification to the complex mechanisms that govern the response to hypoxia, we chose this type of model as it has proven successful in modelling hyperthermia and is suitable for experimental validation. In the proposed model (Eq. 2), there are three compartments representing the fraction of cells at each point in space and time that are in one of three states: alive (*A*), vulnerable (*V*) and dead (*D*), where *A*+ *V* + *D* = 1. We assume that there is a forward pathway both from *A* to *V* and from *V* to *D*, but that there is only a backward pathway from *V* to *A* (i.e., the process from *A* to *V* is reversible, but that from *V* to *D* is irreversible):

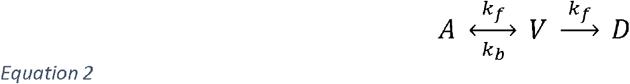

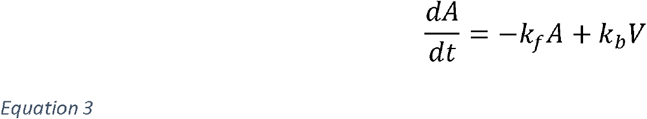

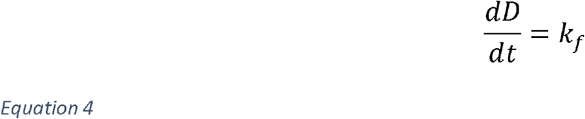

where *k*_*f*_ is the forward and *k*_*b*_ the backward rate constant, both of which are assumed to depend solely on the hypoxic fraction (*H*) in a linear manner:

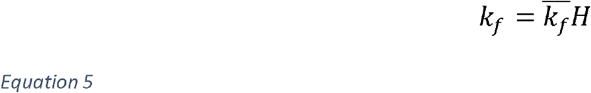

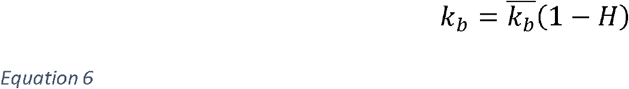

where 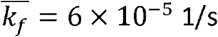 1/s is the forward rate constant and 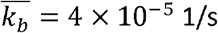 1/s is the backward rate constant, which are forward rate at full hypoxia and backward rate at zero hypoxia respectively. These two constants will be further justified in Section 3.4. Based on the Green’s function simulations, the hypoxic fraction can be expressed as a sigmoidal function of blockage fraction (Section 3.3). The ordinary differential equations (ODEs) were solved by ‘odeint’ function in SciPy.

## 3 Results

### 3.1 Capillary networks and Green’s function method solutions

One example of a 375-micron statistically accurate capillary network cube is shown in Fig. 1a, where the vessel diameter is represented by the line thickness. Figures 1b and 1c display the tissue PO_2_ simulations generated by the Green’s function method inside the same network under normal and hypoxic conditions respectively, where the oxygenation level is indicated by different colours. Figures 1e and 1f show the tissue oxygenation and the positions of intersecting vessels on the centre plane of the same cube (Fig. 1d) The tissue PO_2_ is distributed heterogeneously inside the cube. However, it is closely correlated with the geometry of the microvasculature which can be observed in both 3D and 2D plots.

### 3.2 Independence of oxygen transport on microvascular geometry and length scale

Figure 3 shows the values of average tissue PO_2_ and hypoxic fraction (fraction of tissue with PO_2_ below 10 mmHg) for values of PO_2, in_ in the range 13 to 50 mmHg in the capillary cubes at two sizes (375 μm and 625 μm). The error bar shows the standard deviation of the results among the 10 different cubes at each size, illustrating that the variability between different geometries is relatively small. It can be seen that the average tissue PO_2_ varies approximately linearly with the PO_2, in_ within this range. The hypoxic fraction however has a strongly non-linear relationship with PO_2, in_, with the hypoxic fraction increasing from 20% to 90% when PO_2, in_ varies from 30 to 20 mmHg.

**Figure 3.**
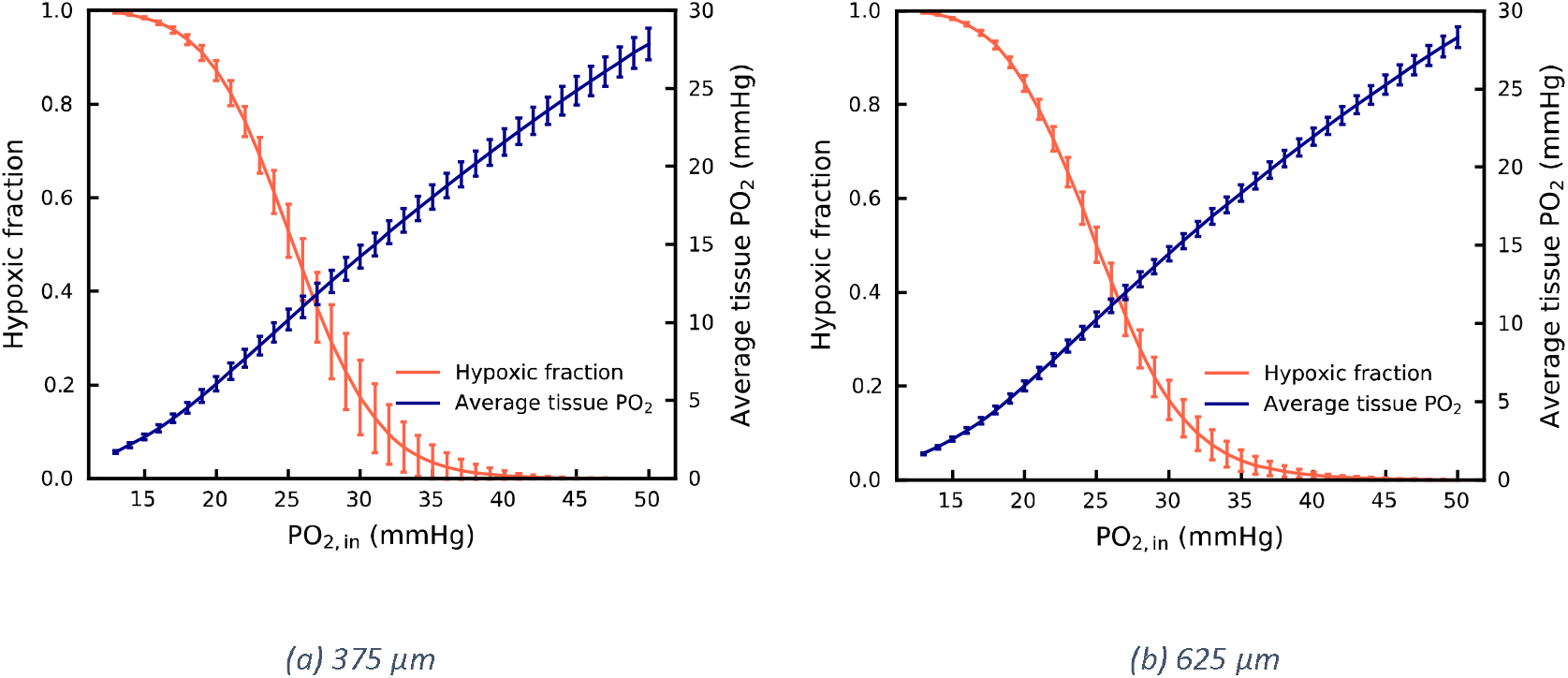
Hypoxic fraction and average tissue PO_2_ in the capillary networks at 375 μm (a) and 625 μm (b). The error bar shows the standard deviation of the results among 10 different cubes.

Figure 4 presents the values of CMRO_2_ and OEF in the same range of PO_2, in_ in the capillary cubes at the two sizes. The error bar also shows the standard deviation of the results among 10 different cubes. When PO_2, in_ changes from 50 to 13 mmHg, the CMRO_2_ displays a decrease from 3 to 0.5 cm^3^/(100cm^3^ · min) and the OEF increases from 0.35 to 0.95 to compensate for the shortage of oxygen supply, both in a strongly non-linear manner.

**Figure 4.**
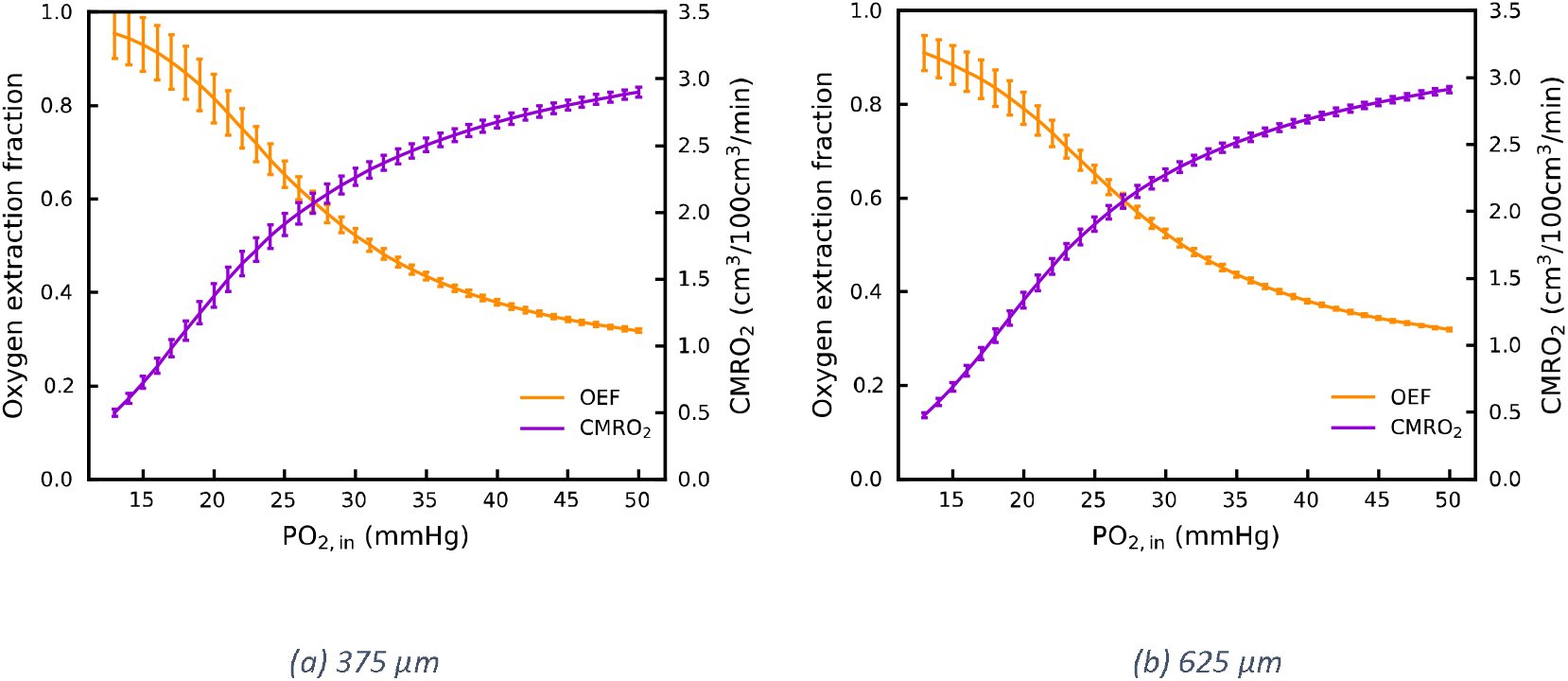
Oxygen extraction fraction and oxygen consumption rate in the capillary networks at 375 μm (a) and 625 μm (b). The error bar shows the standard deviation of the results among 10 different cubes.

The small standard deviation of results between the different capillary cubes indicates that the oxygen transport at the microvasculature scale is largely independent of the specific geometry of the individual capillary networks, as long as the geometrical properties of the networks are maintained. The standard deviation of the results in the 625-micron cubes is smaller than in the 375-micron cubes, which indicates that the effect of microvascular topology and geometry on the oxygen transport becomes increasingly negligible in a larger simulation domain. There is a very close match between the results in the cubes in two different sizes (Figs. 3a and 3b; Figs. 4a and 4b) suggesting that our oxygen transport simulation is length scale independent at a length scale above a few hundreds of microns, which is in very good agreement with the results for permeability^15^. This independence of length scale enables us to use our simulation results directly in representing oxygen transport to the human cerebral microvasculature at longer length scales.

### 3.3 Microthrombi have significant impact on tissue hypoxia beyond 10% vessel occlusions

Figure 5 presents the change in perfusion and hypoxic fraction when the vessels in the capillary network are blocked up to a 20% vessel fraction to mimic the presence of micro-thrombi. In Figure 5a, the perfusion is found to decrease linearly with the increase of blockage fraction in this range, which agrees with the previous simulations^24,26^. The slope of decrease in this study is −3.1 %/%, which is in excellent agreement with the values of − 2.5 %/%^24^ and −3.2 %/%^26^ reported in previous studies. In Figure 5b, the hypoxic fraction remains almost unchanged until the blockage fraction reaches 0.1. Beyond this threshold, the hypoxic fraction is then found to increase steeply with a further increase in blockage fraction. To describe the results of the simulations in a form suitable for larger-scale simulations, we characterized the results for hypoxic fraction, which was fitted to a sigmoidal relationship dependent upon the blockage fraction (*B*) as:

**Figure 5.**
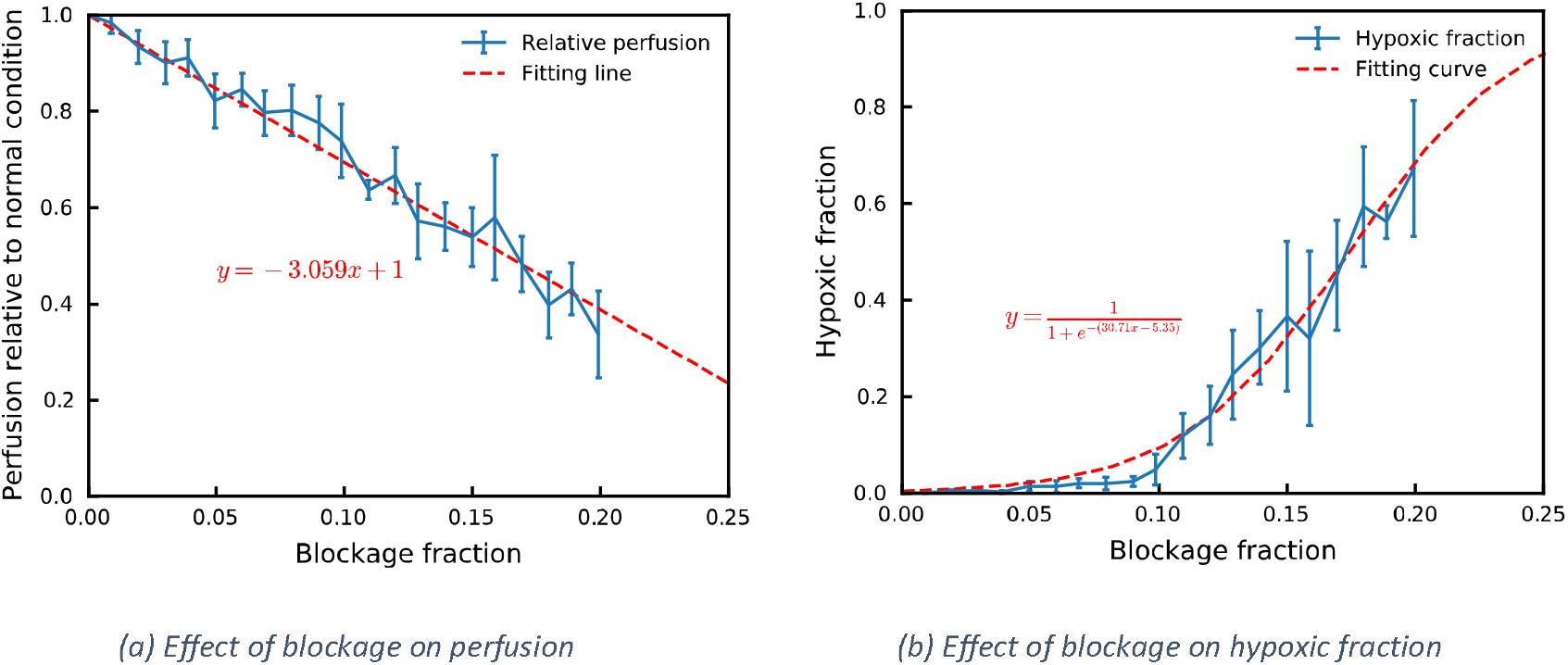
The perfusion (a) and hypoxic fraction (b) as functions of microthrombi blockage fraction.

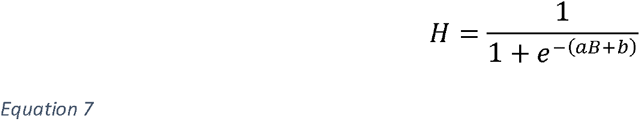

where a = 30.71 and b = −5.35 are two constants derived from least-squares error curve fitting in SciPy. Note that we chose the relationship above since this ensures that the hypoxic fraction remains in the interval 0 to 1.

Note that at a 10% blockage fraction, there are 56 microthrombi inside a 375-micron capillary cube with a total volume of 8.43 × 10^−9^ mL, assuming that there is one thrombus per vessel and that each thrombus has a volume equal to that of a sphere with the same diameter as the blocked vessel. This results in a microthrombi volume of 1.60 × 10^−7^ mL per mL cerebral microvascular volume. If we assume that there are micro-embolisms in a territory of 100 mL fed by middle cerebral artery, the total microthrombi volume will be 1.60 × 10^−5^ mL. Considering the volume of a thrombus is 0.17 mL^38^ and 25% of clot fragments are transported into capillary networks, the total microthrombi volume will be 4.25 × 10^−2^ mL, which is about 2,600 times more than the volume predicted from blocked vessels. These results indicate that the microthrombi in larger vessels must be considered to match the thrombus volume, due to the third power of radius in sphere volume. The results of volume here thus seem to provide a lower limit of the thrombi volume required to give a 10% micro-embolism blockage fraction.

### 3.4 Cell viability with increasing micro-embolism

Figure 6 shows the cell viability predicted from the proposed 3-state cell death model under a range of scenarios. In these 6 example predictions, the initial blockage fraction ranges from 0 to 25% with a step of 5% in order to demonstrate the model response under different conditions. When there is no occlusion in the capillary network (Fig. 6a), the cell viability maintains near 100%. The small gap between alive cell fraction and 100% is negligible over the time scale of usual clinical practice.

**Figure 6.**
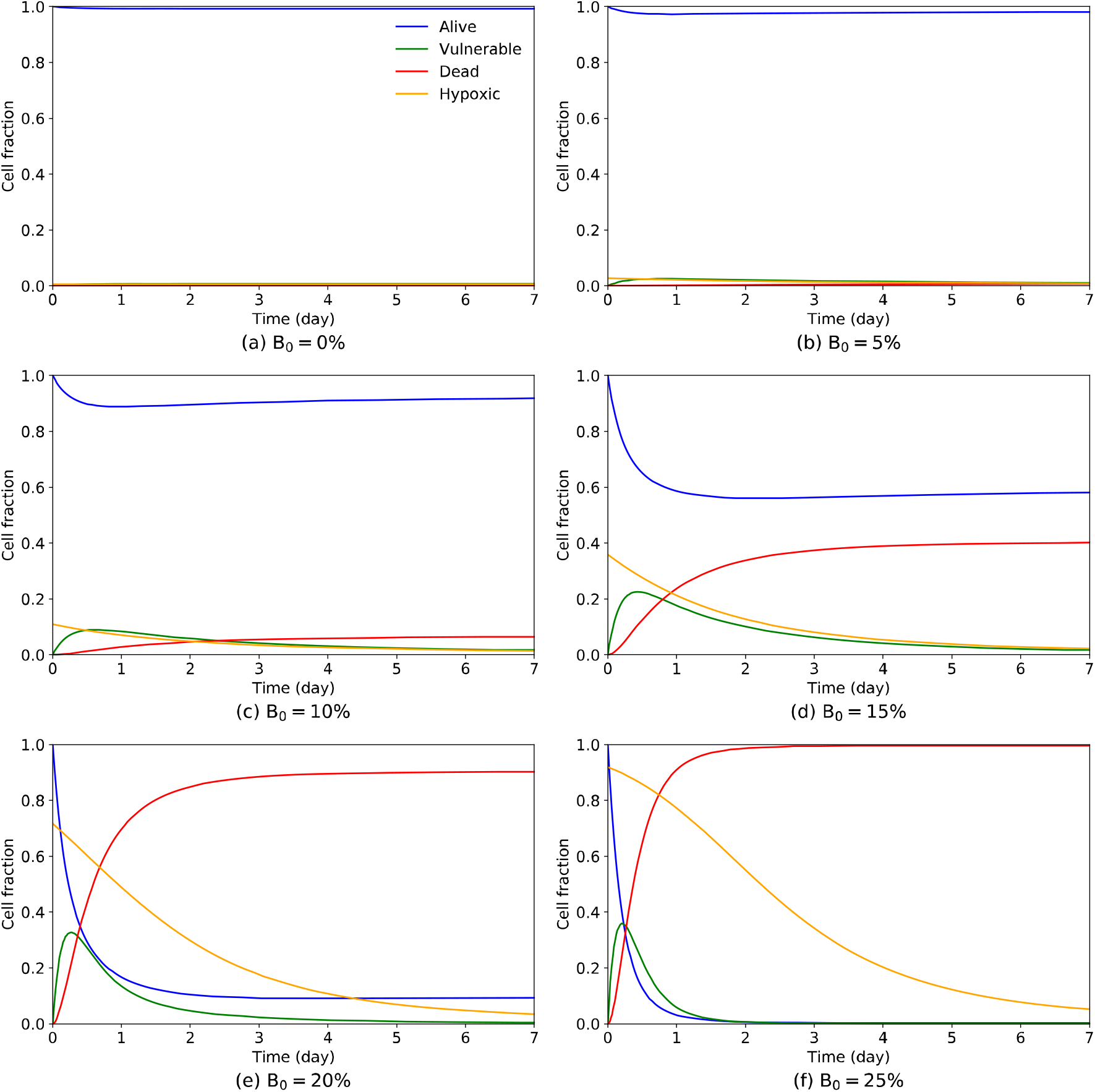
Predicted cell viability and hypoxic fraction over 7 days with different initial microthrombi blockage fraction.

With a small initial blockage fraction of 5% (Fig. 6b), most of the cells still remain alive as shown by the small hypoxic fraction. This can be well explained by the interconnected nature of the topology of the cerebral microvasculature, which gives good resistance to mild microthrombi occlusions. As the blockage fraction increases from 10% to 20% (Figs. 6c, 6d and 6e), however, there is a very substantial change in both dynamic response and steady state value of the model behaviour. The sharp increase in cell death speed and final cell death fraction is caused by the sigmoidal dependence of hypoxic fraction with blockage fraction, which is highly non-linear (Fig. 5b). The blockage fraction of 10% thus appears to be a threshold deciding the final cell death fraction. Beyond the threshold, a further increase in blockage fraction contributes little to the model behaviour (Fig. 6f), since most of the tissue regions have become hypoxic (below 10 mmHg, based on our definition).

Figure 7 shows a sensitivity analysis on the forward rate constant 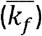 and the backward rate constant 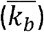 in the cell death model, which are currently unknown and will be fitted with the experimental data in a future study. The final dead fraction at the end of Day 7 with forward and backward rate constants in the range 1× 10^−5^ to 1× 10^−3^ 1/s is plotted for a range of different initial blockage fractions. The combination of rate constants selected for use in this paper is indicated with the red cross. The clear and narrow boundary between the yellow region (high dead fraction) and the blue region (low dead fraction) indicates that there is a distinct threshold for the combination of two rate constants when simulating the final tissue damage. This threshold can be explained by the vessel extravasation model (Eq. 1) where there is no tissue damage when the forward death rate, which is proportional to the forward rate constant and hypoxic fraction, is much smaller than the extravasation rate under a certain initial blockage fraction. The threshold can also be explained by the relationship between the two time constants. When there is a much larger backward rate than forward rate, there is also no tissue damage due to many more cells being saved from the vulnerable compartment than dying. This can be observed as the blue regions at the top left corner of each plot in Fig. 7. These all contribute to the highly non-linear behaviour of the cell death model.

**Figure 7.**
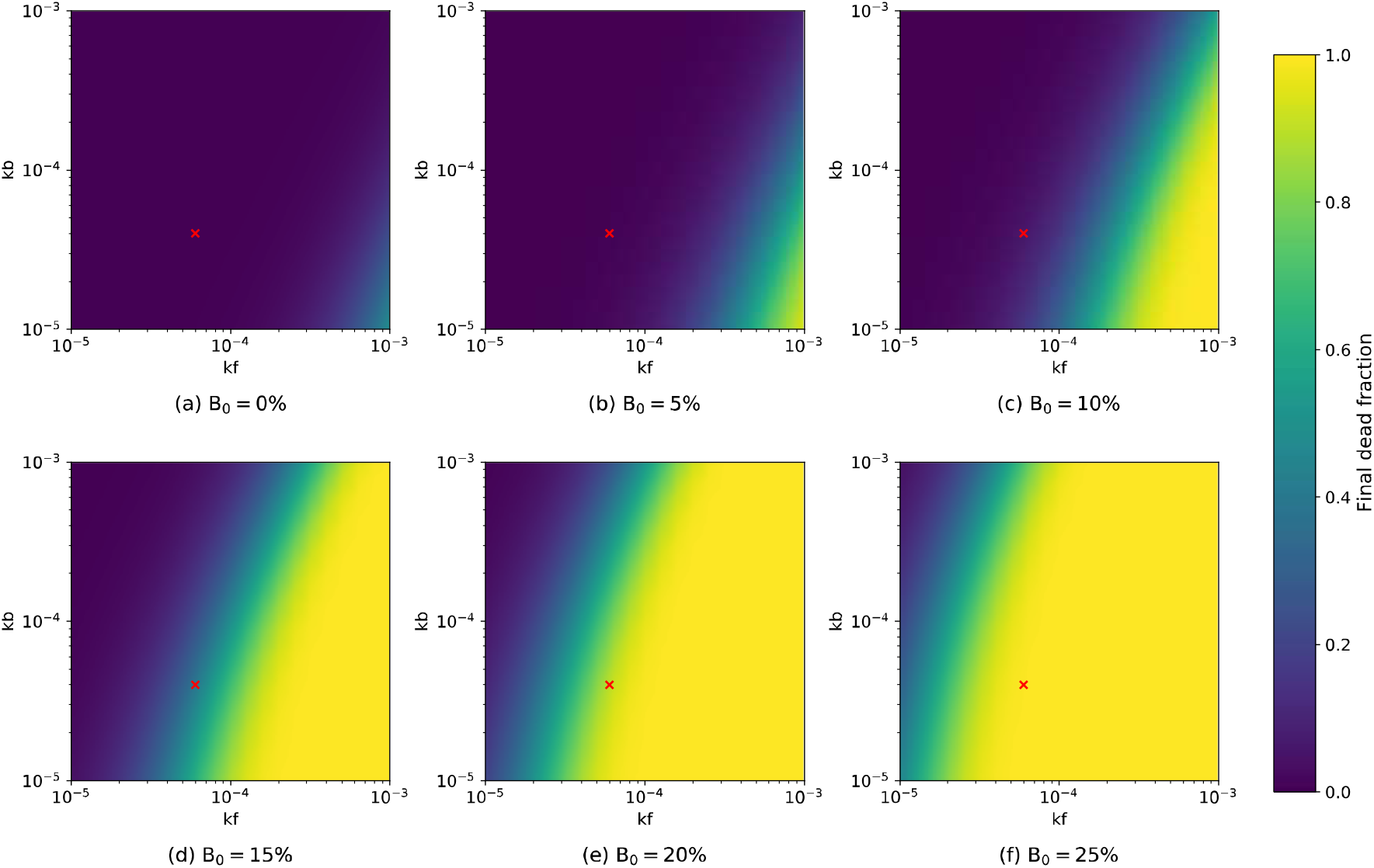
Final dead fraction at the end of Day 7 under different initial blockage fractions and combination of rate constants in the 3-state cell death model. The forward and backward rate constants used in the paper are indicated as the red cross.

It should finally be noted that the cell death simulation here moves from the local behaviour inside a capillary cube towards ODEs at the global scale, based on the independence of oxygen transport on the microvascular geometry and length scale (Section 3.2). It guarantees that the cell death model can be used in whole-brain models easily.

## 4 Discussion

In this paper, we present a new way to quantify and to analyse the effect of cerebral microthrombi on oxygen transport and tissue viability. To our knowledge, this is the first numerical study that investigates the effects of microthrombi on tissue hypoxia. The oxygen transport inside the cerebral microvasculature is strongly influenced by the network geometry and thus leads to a heterogeneous distribution of oxygen levels. However, the overall feature of cerebral oxygen transport in a representative elementary cube is found to be essentially largely independent of the microvascular geometry, which is in line with our previous study on cerebral blood flow^15^. In addition, the oxygen transport results are found to be independent of length scale. The vessel blockage fraction has a linear impact on perfusion, which agrees with previous simulations^24,26^. However, the relationship between blockage fraction and hypoxic fraction is found to be strongly non-linear, modelled here with a sigmoidal function. With the extravasation model and the cell death model, the impact of a certain number of microthrombi on tissue viability can be predicted at a given time after the formation of such microthrombi.

The periodic, statistically accurate microvasculature cubes used here remove the need to reconstruct the vessel geometries in detail and to set the boundary conditions. The cubes can also be connected to represent a larger microvascular domain due to its periodicity, and this advantage has been demonstrated in recent studies^20,39^. The Green’s function method significantly reduces the computational costs of solving the oxygen transport in a complex network by transforming 3D diffusion equations into a 1D Green’s function. This study implements the Green’s function method in statistically accurate capillary networks based on human morphometric data^17^, which provides a pathway to investigate oxygen transport in the human microvasculature with limited experimental data and much reduced computing power.

The prediction of irreversible tissue damage is always a tricky task, which is usually neglected in cerebral metabolism models^40–42^. Here we implemented a 3-state cell death model, a similar form of which was used in a previous hyperthermic cell death study. The model has the advantage of modelling the various cell death rate under different conditions. In this study, temperature-related rate constants in the 3-state model are replaced with hypoxia-related rate constants so that the cell death rate is positively correlated with tissue hypoxic fraction and the forward pathway will shut down as hypoxia is relieved by the extravasation of microthrombi^12,28,29^.

The primary advantage of this workflow is its possibility to scale up the geometry of the cerebral microvasculature, its function in oxygen delivery and hypoxic damage caused by microthrombi to the tissue scale easily, and to couple with existing models at both a cortical column scale^16,20,26^ and a whole brain scale^22^. Another advantage is its ability to locate the hypoxic regions and microinfarcts and to relate them to the distribution of microthrombi. This makes it possible to be validated against *in vivo* micro-embolism experiments that cannot be easily done with computational fluid dynamics or agent-based models.

One limitation of this study is the assumption of no oxygen supply from larger vessels like arterioles. However, the oxygen transport from vessels to tissue happens over several vessel generations^36,43^. The oxygen transport from larger vessels cannot be simulated with the current capillary cubes, which limits the length scale at which this model can be used. In future studies, the oxygen transport from larger vessels could be modelled when the microvascular model is coupled with the penetrating vessel models^16^, as has previously been performed with blood flow^20^.

The second limitation is the partial use of animal data and parameters, when this study is intended to be applied on human cerebral microvasculature. Rat data have been used to derive the parameters in the Hill equation for blood oxygen saturation used in the Green’s function method^27^ and the data used for fitting the extravasation model^12^. Here we assume that these physiological relationships or processes are very similar in human and rats. However, once more human data are available, these models can easily be updated.

The models proposed in this study are purely passive, although local perfusion is actively and tightly controlled. The control mechanisms are very complex and often include other biomarkers like pH, carbon dioxide and nitric oxide and cells like pericytes and astrocytes, which are beyond the scope of this study^32^, although this will be the subject of future research.

Lastly, this paper focuses on the averaged properties inside the capillary cubes. Due to the heterogeneous distribution of tissue oxygenation (Fig. 1), there might be concentrated hypoxic regions even with a low hypoxic fraction. The distribution of localised hypoxic regions will be further investigated and potentially validated in our future study.

In summary, this paper has applied the Green’s function method on periodic, statistically accurate human capillary networks to investigate the oxygen transport inside the cerebral microvasculature. With a novel extravasation model and a novel 3-state cell death model, the effect of microthrombi on oxygen transport and hypoxic cell death can be simulated. Such an approach will be a key part to the prediction of tissue damage in response to microthrombi.

## Acknowledgement

This work was partially funded by the European Union’s Horizon 2020 research and innovation programme, the INSIST project, under grant agreement No 777072.

## Appendix A Governing equations for blood flow

In general, blood flow is governed by the Navier-Stokes equations. Under the assumption of steady-state, axisymmetric and fully-developed flow along the direction of the blood vessels, the blood flow inside the capillary network can be represented by Hagen-Poiseuille equation:

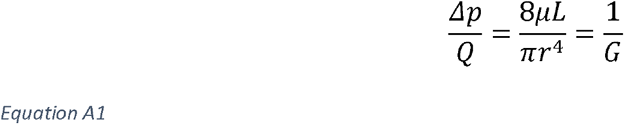

where *p* is pressure, *Q* is flow rate, *L* is the vessel length, *r* is the vessel radius, *G* is the flow conductance and µ is the apparent blood viscosity, which is an empirical equation of vessel diameter and haematocrit^44^.

Since there is a linear relationship between pressure drop and flow rate in each vessel segment, the flow characteristics inside the capillary network can be described by the conductance matrix as:

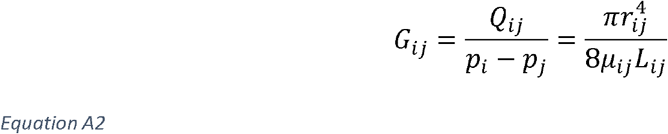

where *i* and *j* are boundary nodes of capillary segments and *Q*_*ij*_ is the flow rate from *i* to *j. G*_*ij*_ is equal to 0, if there is no connection between node *i* and node *j*. Due to flow conservation at each node, the pressure at each capillary segment node inside the network can be solved as:

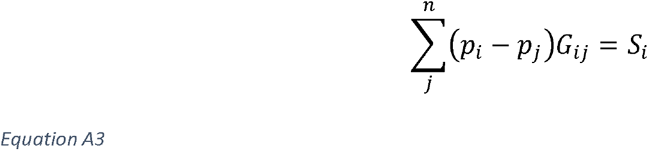

where *S*_*i*_ is the source term at each node. The flow in each segment can thus be solved using Eq. A2.

## Appendix B Governing equations for Green’s function method and oxygen reaction-diffusion models

The Green’s function method has been introduced in detail in previous studies^27,33^. The oxygen transport in brain tissue can be simplified as a reaction-diffusion equation:

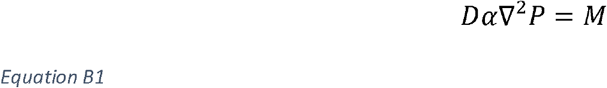

where *D* is the diffusion coefficient, α is the tissue oxygen solubility and *P* is the partial pressure of oxygen and *M* is the metabolic rate. According to potential theory, the steady-state diffusion of oxygen is rewritten as a Green’s function between every point source of capillary segments and every point sink evenly distributed in the tissue as:

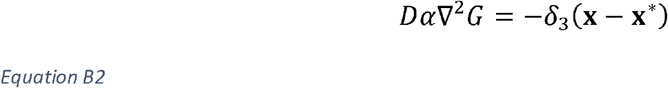

where *G* is the Green’s function, δ_3_ is a three-dimensional Dirac delta function, **x** = (*x*_1_, *x*_2_,*x*_3_) is a point inside the tissue and 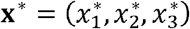 is a point source on the capillary surface. In an infinite domain, the Green’s function can be solved as:

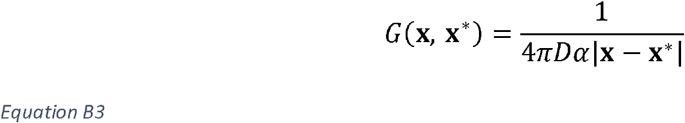

By integrating the oxygen potentials from all sources (surfaces of all capillary segments), the oxygenation of each tissue point can be calculated as:

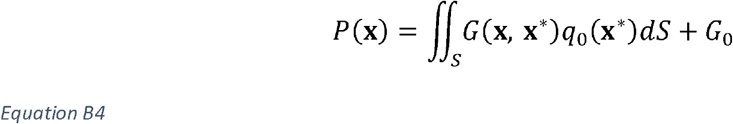

where *S* is the total tissue-vessel boundaries, *q*_0_(**x**\*) is the distribution of the source strengths on these boundaries and *G*_0_ is a constant updated in each iteration.

In this study, 25 tissue points were placed along each dimension of the capillary cubes to give good spatial resolution (Figs. 1). The spacings of tissue points are 15 μm (375 μm cubes) and 25 μm (625 μm cubes), which match the range of 10-30 μm used in a previous study^27^. The maximum capillary vessel segment length required by the algorithm was set as 100 microns, which ensured that any vessel segment longer than 100 microns would be divided into smaller segments.

The consumption of each tissue point is represented by a Michaelis-Menten relationship with oxygenation:

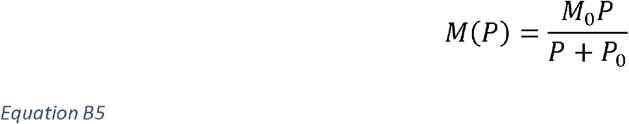

where *M*_0_ is the maximum metabolic rate of oxygen and *P*_0_ is the Michaelis-Menten constant.

The oxygen transport in blood vessels follow the 1D mass transport equation. The blood oxygen concentration can be expressed as:

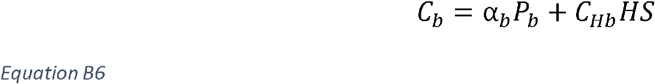

where *α*_b_ is the blood oxygen solubility, *P*_*b*_ is blood PO_2_, *C*_*Hb*_ is the haemoglobin-bound oxygen capacity in fully saturated red blood cells, H is the haematocrit and S is the oxygen saturation of haemoglobin. The oxygen saturation of haemoglobin follows a Hill equation with partial pressure of oxygen as:

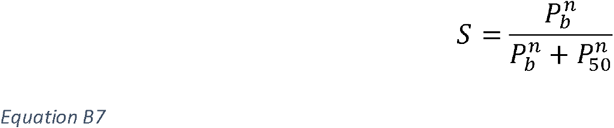

where *n* is the Hill equation exponent and P_50_ is PO_2_ at half maximal haemoglobin saturation. All the Green’s function parameters are the same as previously used^27^, except for values of maximum metabolic rate of oxygen and haematocrit that are quoted in the main text.

## Footnotes

1 https://physiology.arizona.edu/people/secomb/greens

2 https://scipy.org

